# Human stem cell derived sensory neurons are positioned to support varicella zoster virus latency

**DOI:** 10.1101/2020.01.24.919290

**Authors:** Tomohiko Sadaoka, Labchan Rajbhandari, Priya Shukla, Balaji Jagdish, Hojae Lee, Gabsang Lee, Arun Venkatesan

## Abstract

The neuropathogenesis of varicella-zoster virus (VZV) has been challenging to study due to the strict human tropism of the virus and the resultant difficulties in establishing tractable experimental models. *In vivo*, sensory neurons of the dorsal root ganglia and trigeminal ganglia serve as cellular niches that support viral latency, and VZV can subsequently reactivate from these cells to cause disease. Whether sensory neurons possess intrinsic properties that position them to serve as a reservoir of viral latency remains unknown. Here, we utilize a robust human sensory neuron system to investigate lytic infection and viral latency. We find that sensory neurons exhibit resistance to lytic infection by VZV. On the other hand, latent infection in sensory neurons is associated with an episomal-like configuration of viral DNA and expression of the VZV latency-associated transcript (VLT), thus closely mirroring the *in vivo* state. Moreover, despite the relative restriction in lytic infection, we demonstrate that viral reactivation is possible from latently infected sensory neurons. Taken together, our data suggest that human sensory neurons possess intrinsic properties that serve to facilitate their role as a latent reservoir of VZV.

**IMPORTANCE:** Varicella-zoster virus (VZV) has infected over 90% of people worldwide. Following primary infection, the virus can remain dormant in the nervous system and may reactivate later in life, with potentially severe consequences. Here, we develop a model of VZV infection in human sensory neurons in order to determine whether these cells are intrinsically positioned to support latency and reactivation. We find that human sensory neurons are relatively resistant to lytic infection, but can support latency and reactivation. Moreover, during *in vitro* latency human sensory neurons, but not other neurons, express the newly discovered VZV latency-associated transcript (VLT), thus closely mirroring the *in vivo* latent state. Taken together, these data indicate that human sensory neurons are uniquely positioned to support latency. We anticipate that this human sensory neuron model will serve to facilitate further understanding of the mechanisms of VZV latency and reactivation.

## INTRODUCTION

Varicella-zoster virus (VZV) is a neurotropic human alphaherpesvirus that has infected over 90% of people worldwide. Following primary infection, the virus establishes lifelong latency in sensory neurons of the cranial nerve and dorsal root ganglia (DRG) with the potential for reactivation later in life (1, 2). Viral reactivation can have severe consequences, including herpes zoster (shingles), encephalitis, and myelitis (3, 4), and has recently been associated with giant cell arteritis (5). Worldwide, an increasingly aging population and the more widespread adoption of novel immunosuppressive therapies for autoimmune conditions place individuals at growing risk for viral reactivation (6), pointing to the need to better understand the neuropathogenesis of VZV.

The cell type specificity of VZV latency remains poorly understood, in part due to lack of tractable models to study viral neuropathogenesis. Primary dissociated neurons from adult human sensory ganglia have been isolated and infected, as have fetal DRG (7, 8). These models have been useful in demonstrating some aspects of VZV-neuronal interactions; for example, in SCID (severe combined immunodeficiency) mice xenografted with human fetal DRG, viral replication occurred in a subset of neurons but was blocked in cells that expressed the mechanoreceptor marker RT97 (9–11). More recently, *in vitro* neuronal models of VZV infection utilizing human neural stem cells (12) and human embryonic stem cells (ESC) (13) have been developed. We and others have used such models to study various aspects of VZV neuropathogenesis, including characterization of transcriptional changes in infected cells, demonstration of viral axonal transport, exploration of the role of the cellular JNK pathway in viral infection, and establishment of models of viral latency and reactivation (13–18).

Despite these advances, a tractable model of human sensory neuronal infection by VZV has, to date, been elusive (19). As a result, it has remained unclear as to whether sensory neurons possess intrinsic features that contribute to their role in VZV neuropathogenesis in humans. Here, we utilize a robust human induced pluripotent stem cell (iPSC)-derived sensory neuron system to test the hypothesis that sensory neurons *in vitro* are uniquely poised to support VZV latency and reactivation.

## RESULTS

### Characterization of a human sensory neuron system to study VZV infection

To develop a robust sensory neuron platform for the study of VZV infection, we utilized human sensory neuron progenitor cells to generate mature human sensory neurons (HSNs). By two weeks of differentiation, over 90% of cells expressed the pan-neuronal marker neurofilament (**Fig 1A**). Expression of sensory neuronal markers peripherin and brain-specific homeobox/POU domain protein 3A (Brn3a) was noted in subsets of cells, and occasional pockets of Islet 1 positive cells were also noted. By four weeks of differentiation, Brn3a is clearly localized to the nucleus (arrows, top panel in **Figure 1B**) in Tuj1 (the neuronal cytoplasmic marker beta III tubulin) positive neurons, in keeping with its role as a nuclear homeodomain transcription factor. Peripherin, an intermediate filament protein, is localized to the cytoplasm and neuronal processes, with a similar expression pattern to Tuj1 as expected (middle and bottom panels in **Figure 1B**). Subsets of voltage-gated ion channels, Na_v_1.7 and Na_v_1.8 positive cells, each of which colocalize with Tuj1 and peripherin, are observed (arrows, middle and bottom panels, respectively in **Figure 1B**). By six weeks of differentiation, >90% of cells are positive for peripherin and Brn3a, >80% are positive for Islet 1 and >50% are positive for Na_v_1.7 and Na_v_1.8, indicating a mature sensory neuron phenotype (top panel in **Fig 1C**). In contrast, our human ESC-derived neurons (16, 18) represent a mixed population (human mixed neurons; HMNs) that express CNS (central nervous system) markers such as GABA_A_ receptor, Glycine receptor, and CTIP1 along with robust expression of the pan-neuronal markers Map2 and Tuj1 (16, 18). Expression of the sensory neuronal markers peripherin, Brn3a, Islet 1, and Na_v_1.8 was rarely (<1% of cells) observed in HMNs (lower panel in **Fig 1C**).

**Figure 1.**
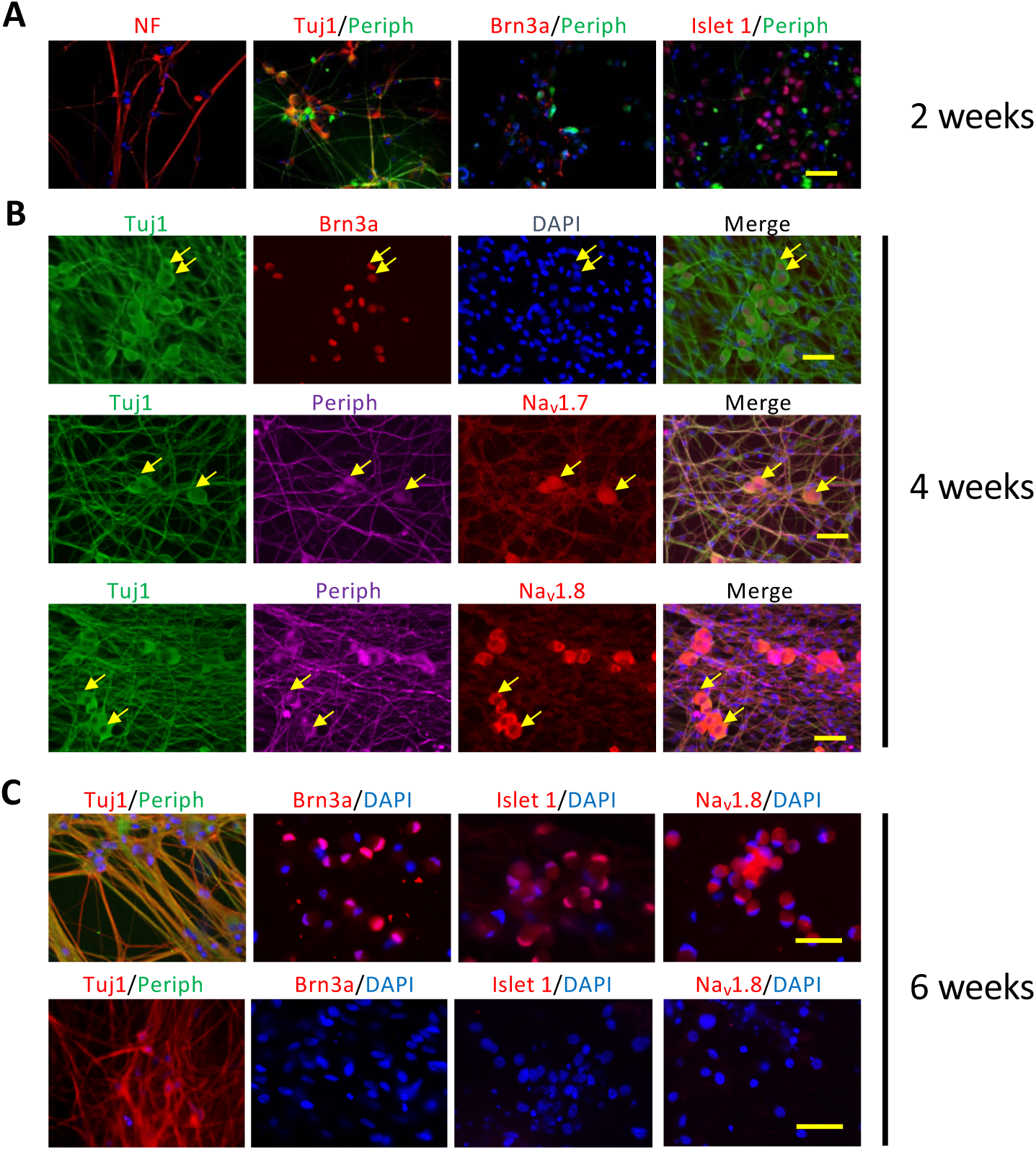
Characterization of a human sensory neuron system to study VZV infection. Human sensory progenitor cells are differentiated over the course of six to seven weeks. At two weeks (A), many cells express the pan-neuronal markers neurofilament (NF) and beta III tubulin (Tuj1). Some cells express peripherin, a marker found mainly on neurons of the peripheral nervous system, and expression is colocalized with Tuj1. Brn3a and Islet 1, markers of sensory neurons, begin to be expressed in subsets of cells. By four to five weeks (B), many neurons (expressing Tuj1) coexpress Brn3a, and expression of the sensory neuron voltage-gated sodium ion channels Na_v_1.7 and Na_v_1.8 are found in subsets of Tuj1+/peripherin+ sensory neurons. By six to seven weeks (C, upper row), virtually all cells are Tuj1+/peripherin+ sensory neurons. By this time, Brn3a and Islet 1 are appropriately localized to the nucleus, and Na_v_1.8 expression is seen in many cells. In contrast, HMNs (C, lower row), while robustly expressing the pan-neuronal marker Tuj1, are not observed to express Brn3a, Islet 1, or Na_v_1.8. Scale bars; 50 µm.

### Lytic infection of sensory neurons

Following characterization of HSN, we first attempted to infect them with VZV via standard conditions (16, 18). HSNs and HMNs differentiated for six to seven weeks were exposed to cell-free rVZV_LUC_BAC (20), which contains a GFP cassette that is expressed upon infection. While GFP expression was noted in HMNs by day 3 and was robust by day 5, no such expression was observed in HSNs (**Fig 2A**). Since flow cytometry may enable more sensitive detection of GFP, we dissociated infected cells and assessed for GFP expression in a quantitative manner. While 5-12% of HMNs expressed GFP by day 5 following infection as judged by flow cytometry, GFP expression was undetectable in two of three HSN cultures and in the third less than 1% of cells expressed GFP (**Fig 2B**). In addition, while the VZV glycoprotein E (gE) was readily detectable by Western blot from lysates of infected HMNs, no such expression was observed in HSNs (**Fig 2C**).

**Figure 2.**
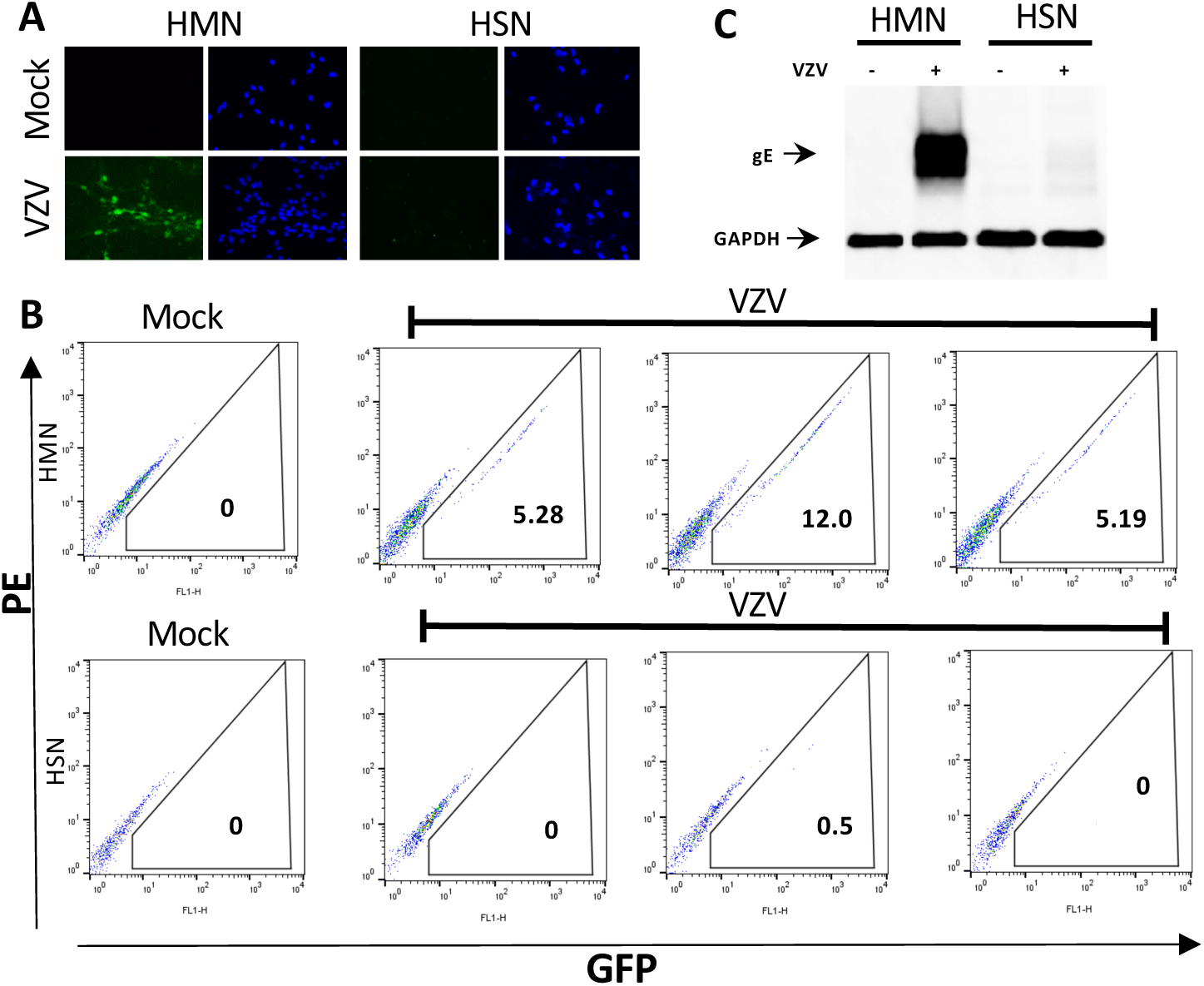
Human sensory neurons are resistant to lytic infection by VZV under standard conditions. (A) Neurons differentiated for six to seven weeks are either mock-infected (Mock) or infected by 100 PFU of rVZV_LUC_BAC (VZV) and observed for GFP expression over five days. Clusters of GFP+ cells are seen in HMNs but not HSNs (B) Flow cytometry confirms that while HMNs robustly support GFP expression, HSNs do not (three separate infections for each condition are shown). GFP channel is plotted on X-axis, while PE channel (negative control) is plotted on the Y-axis. Numbers refer to percentage of GFP+ cells. (C) Western blot analysis of expression of VZV glycoprotein E (gE) in HMNs and HSNs infected by VZV.

### Relative resistance to lytic infection in sensory neurons

Since we occasionally observed limited numbers of HSNs expressing GFP by flow cytometry following infection with rVZV_LUC_BAC under standard conditions (**Fig 2B**), we next determined whether we could establish conditions in which lytic infection in HSNs was more robust. We found that doubling the amount of virus added to HMNs resulted in increased GFP expression, such that 19-26% of HMNs expressed GFP at five days post infection **(Fig 3A, top row; compare to** **Figure 2B, top row**). However, even when using ten times the amount of virus, GFP expression was undetectable in two of three HSN cultures and in the third only 0.14% of cells were found to express GFP (**Fig 3A**). We next sought to determine whether viral nucleic acid was present in HSNs, which might occur despite absence of robust GFP expression, the latter of which depends upon substantial viral replication. We examined transcription of immediate-early (IE) ORF61, early (E) ORF29, and late (L) ORF14 gene. We detected all kinetic classes of transcripts in HMNs differentiated for either four or six weeks and in one set each of the HSNs, though at lower levels (**Fig 3B**). Notably, however, in two of three HSN cultures differentiated for either four or six weeks, only very low levels of ORF61 were detectable while ORF29 and ORF14 were undetectable. We also examined viral DNA and found that HSNs harbored lower levels of VZV DNA as compared to HMNs, regardless of whether neurons were differentiated for four or six weeks prior to infection (**Fig 3C**).

**Figure 3.**
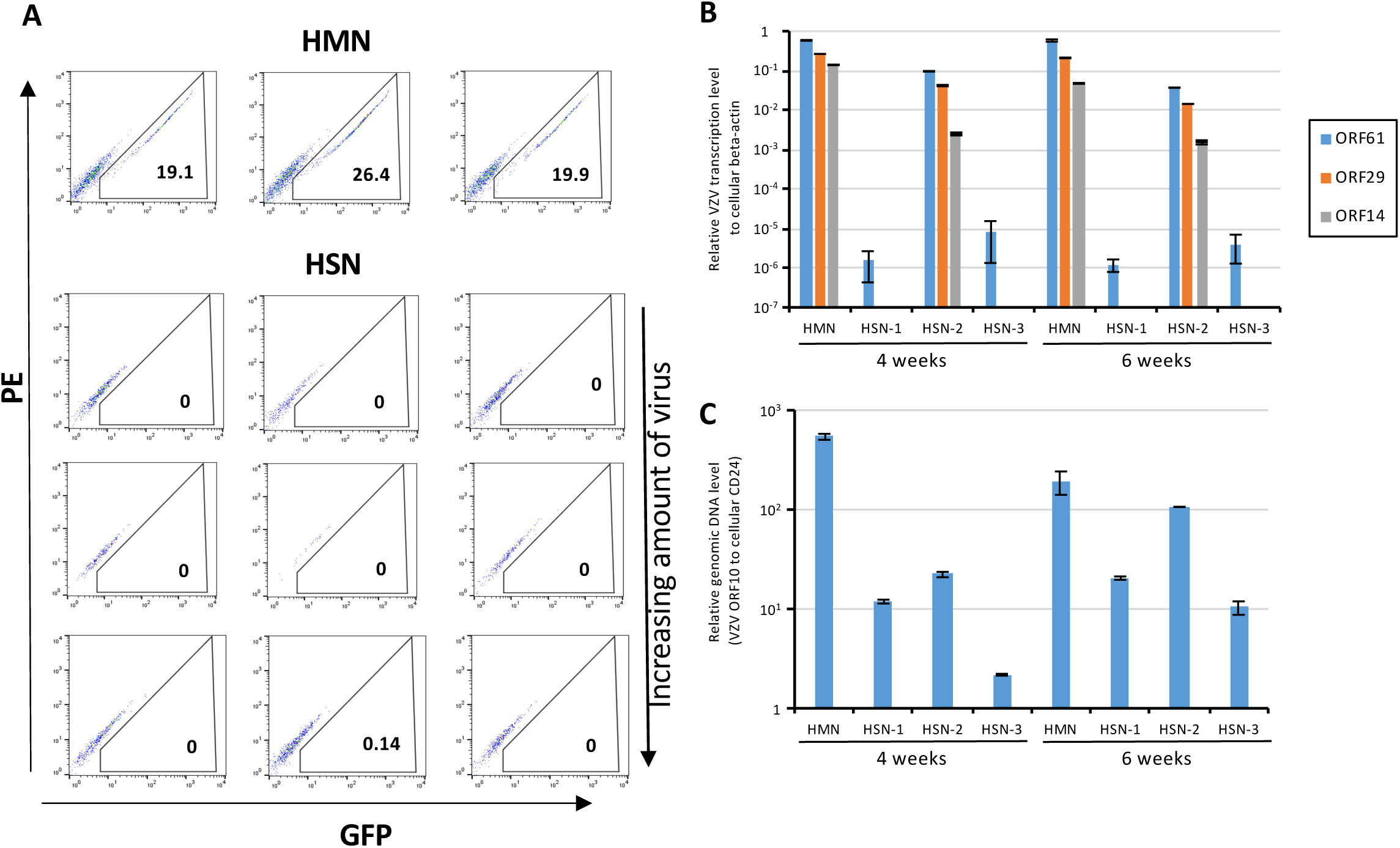
Sensory neurons are relatively resistant to lytic infection. (A) Neurons differentiated for six to seven weeks are infected by rVZV_LUC_BAC and undergo flow cytometry for detection of GFP expression at 5 d.p.i. HMNs are infected with 200 PFU, while HSNs are infected with 400, 800, and 2,000 PFU of virus. GFP channel is plotted on X-axis, while PE channel (negative control) is plotted on the Y-axis. Numbers refer to percentage of GFP+ cells. (B) Transcriptional analysis of infected HMNs and three separate cultures of HSNs (labeled 1-3). (C) qPCR demonstrates that VZV DNA can be detected in each of the HSN cultures, though at substantially lower levels than in HMNs.

We then examined whether the restriction in lytic infection in HSNs is so severe as to preclude the development of productive virus altogether. HSNs differentiated for six weeks were infected for one to two weeks - longer than the standard three to five days - and cells were analyzed for viral nucleic acid. We were able to detect IE, E, and L transcripts at one week following infection, and levels of each had increased by two weeks following infection (**Fig 4A**). Similarly, viral DNA was detected at 1 week post infection (w.p.i.), with increasing amounts observed at 2 w.p.i. (**Fig 4B**). In order to determine whether productive virus was formed from infected HSNs, cells were scraped, placed atop a monolayer of ARPE19 cells and cultured for seven days. Infectious focus formation was confirmed on ARPE-19 cells, and consistent with increasing amount of viral transcripts and viral DNA replication, infectious focus counts were increased from one week to two weeks (**Fig 4C**).

**Figure 4.**
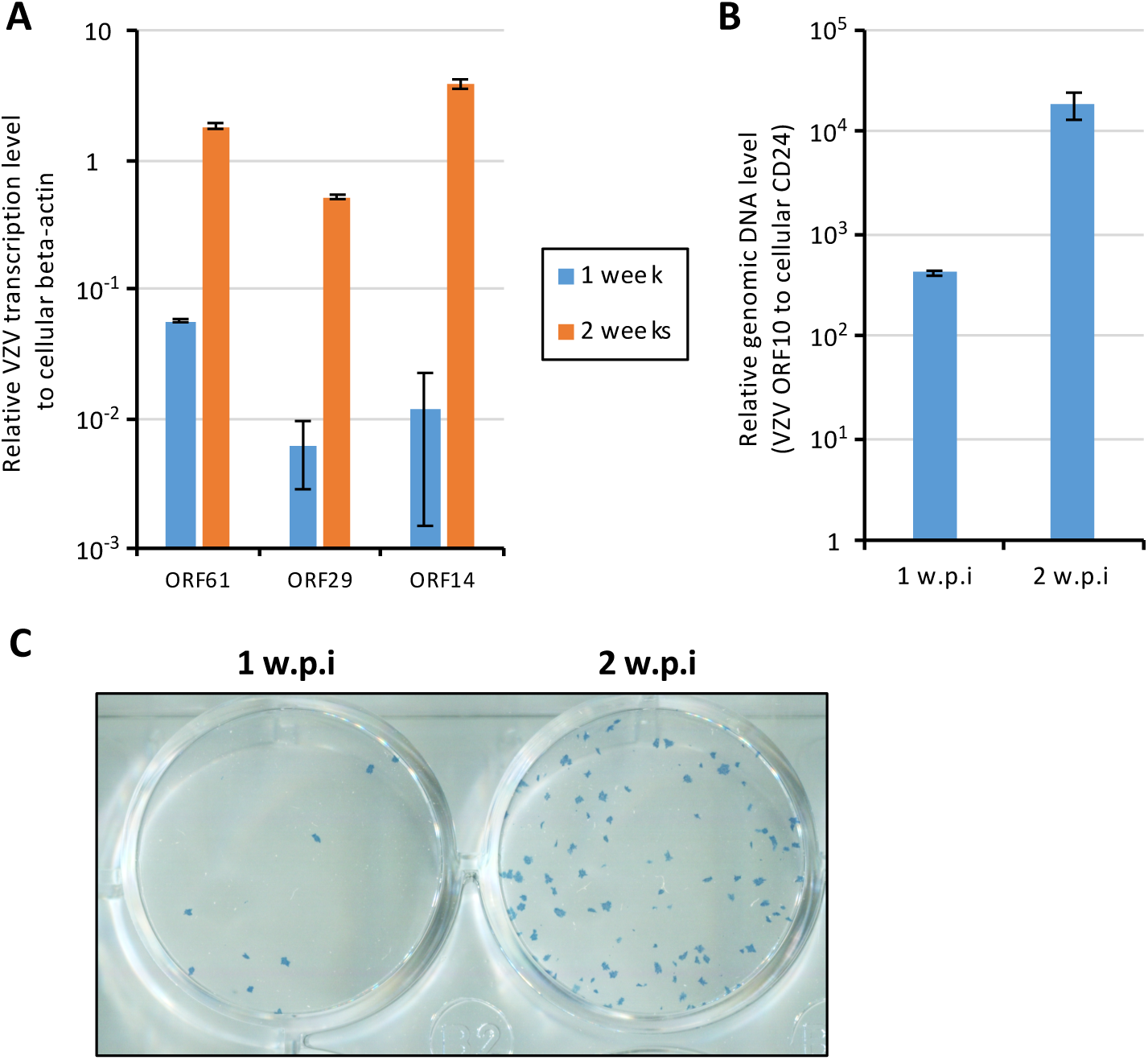
Sensory neurons are capable of supporting productive viral infection. (A, B) HSNs differentiated for six weeks are infected with 400 PFU of pOka VZV for one to two weeks prior to analysis. (A) Transcriptional analysis demonstrates that ORF61, ORF29, and ORF14 are detected at increasing levels from one to two w.p.i. (B) VZV DNA is detected at increasing amounts from one to two w.p.i. (C) Infectious focus forming assay performed from HSNs infected for one or two weeks prior to application atop a monolayer of ARPE19 cells.

### Human sensory neurons are permissive to VZV latency and reactivation *in vitro*

We next examined the establishment of latency in HSNs. We took advantage of a microfluidic platform that we have previously developed that allows for axonal infection of neurons, a factor that appears critical for developing an *in vitro* latent state (16, 18) (**Fig 5A**). In this platform, neuronal cell bodies are cultured in the somal compartment and allowed to extend axons into the fluidically restricted axonal compartment. Cell-free VZV is then added specifically to the axonal compartment, following which an *in vitro* latent state may be established in a small number of neurons (orange cells, middle panel of **Fig 5A**). RT-qPCR using RNA isolated from the somal compartment revealed the presence of the VLT transcript in HSNs via two different primer sets, while VLT was undetectable in HMNs (**Fig 5B**). A primer set targeting ORF63, whose transcript was also detected in human trigeminal ganglia (TG) harboring VZV DNA though at substantially lower levels than VLT (21), did not result in detectable expression in either HSNs or HMNs. We also examined the configuration of the VZV viral genome by using the ratio of terminal repeat joint to genomic linear region abundance, and found the ratios in both HSNs and HMNs to be close to one (**Fig 5C**), indicating a predominantly circular conformation of viral DNA as would be expected were the virus in an episomal configuration as occurs during latency either *in vivo* or *in vitro* (16, 22).

**Figure 5.**
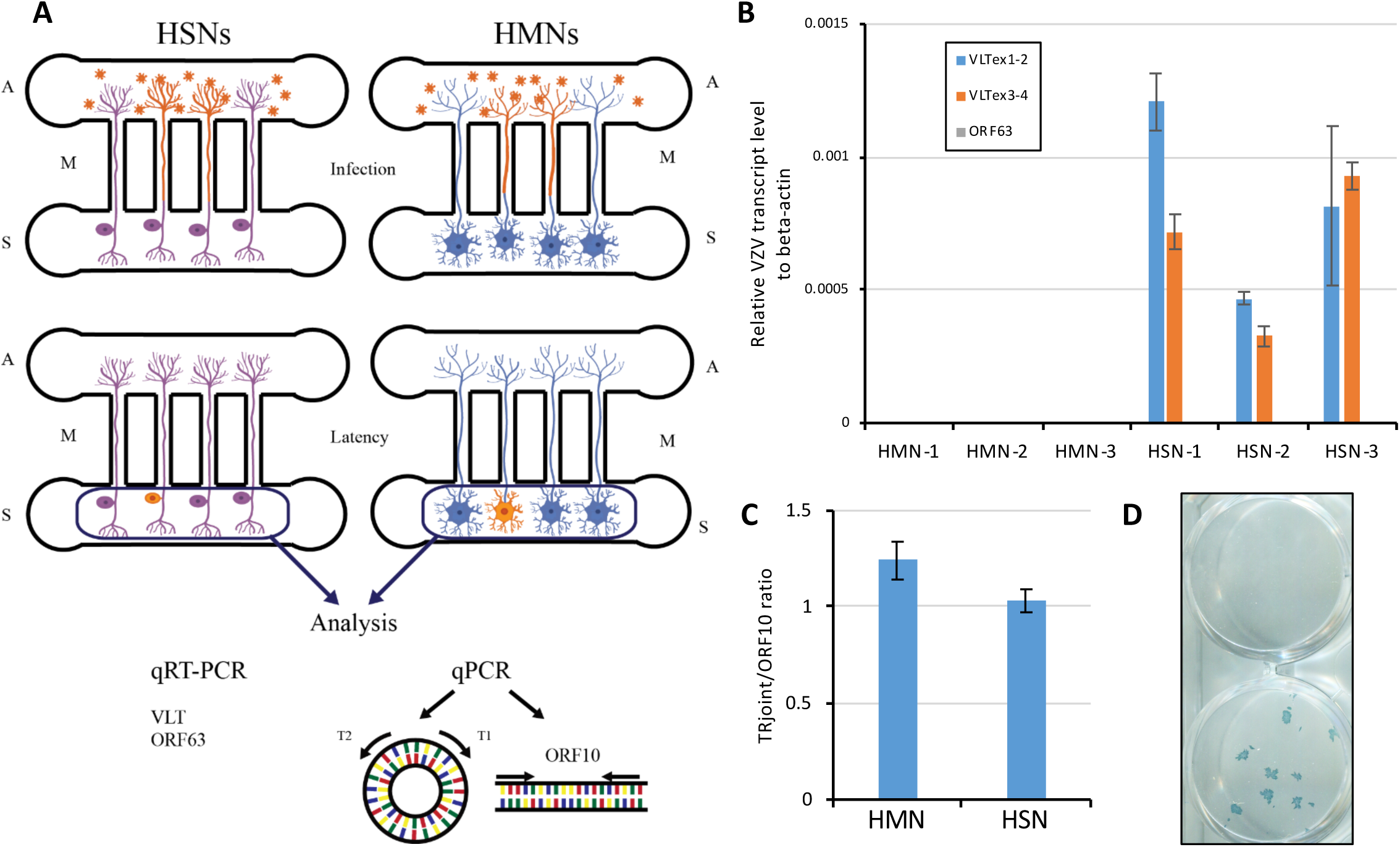
*In vitro* latency in sensory neurons resembles the *in vivo* state. (A) Schematic of *in vitro* latency design. S; somal compartment, A; axonal compartment, M; microchannels. VZV virion and infected neurons are shown in orange. (B) Transcriptional analysis in axonally-infected HMNs and HSNs (3 separate cultures each, labeled 1-3). (C) qPCR to determine the configuration of the viral genome using the ratio of terminal repeat joint and genomic linear region (ORF10) abundance (n=3). (D) Latently infected HSNs were treated with anti-NGF Ab (50 mg/mL) for 14 days to determine whether VZV reactivation would occur. Infectious focus forming assay on ARPE19 cells demonstrates successful reactivation in one of two wells depicted.

Finally, we sought to determine whether viral reactivation could occur following the establishment of an *in vitro* latent state of VZV infection in HSNs. HSNs infected with cell-free VZV from the axonal compartment were cultured for 14 days to establish latency, followed by depletion of neurotropic factors (NGF, GDNF, BDNF and NT-3) and treatment with anti-NGF Ab (50 mg/mL) for 14 days. Transfer onto ARPE-19 cells and culture for a subsequent seven days resulted in two of forty independent samples in which complete viral reactivation was detected via infectious focus forming assay on ARPE-19 cells (**Figure 5D**).

## DISCUSSION

In humans, sensory neurons represent a unique cellular niche that supports VZV latency. Here, we have established a robust human sensory neuron model system *in vitro,* and demonstrated that these cells are relatively resistant to lytic infection by the virus. Latent infection of these sensory neurons appears to closely mimic that of human TG *in vivo* (21). Furthermore, we were able to observe productive viral reactivation from the *in vitro* latent state. These data suggest that human sensory neurons possess intrinsic properties that facilitate VZV latency, and that these properties can be recapitulated *in vitro*.

It has long been recognized that viral latency of many neurotropic alphaherpesviruses is associated with the expression of a single or restricted set of LATs (latency-associated transcripts) that map antisense to the gene encoding the ICP0 (infected cell polypeptide 0) of herpes simplex virus and its analogues (23–25). Recent work has demonstrated that latent infection in human TG by VZV, too, is accompanied by such a transcript, termed VZV latency-associated transcript, VLT (21, 26). Deep sequencing of virus nucleotide-enriched RNA from human TGs with short post-mortem interval consistently demonstrated that viral expression is highly restricted to VLT, often accompanied by ORF63 RNA (21). The DRG and TG represent the only sites of confirmed VZV reactivation from latency *in vivo* (27, 28). Little is known, however, about mechanisms of VZV latency and reactivation in sensory neurons, in part because most human stem-cell derived neuronal culture systems are typically comprised of few, if any, neurons expressing markers of sensory neurons (16). Here, we find that while both sensory and mixed neurons support an episomal configuration of VZV genomic DNA following axonal infection in a microfluidic platform, only sensory neurons recapitulate the *in vivo* expression of VLT. While it has been shown that expression of VLT can suppress expression of the viral transactivator ORF61 (21), the precise role of VLT RNA and/or protein in establishment or maintenance of latency remain to be elucidated. Nevertheless, our data indicate that sensory neurons derived from human iPSC closely mirror the biology of VZV in human TGs with respect to expression of VLT.

In order for latency to be established *in vivo,* sensory and autonomic neuronal ganglia, including TG or DRG, must be infected and lytic infection must presumably be suppressed in order to prevent destruction of the neurons that will subsequently harbor latent infection. Whether lytic infection necessarily precedes establishment of latency is unknown. Routes by which VZV infects sensory ganglion neurons *in vivo* remain unclear, though several non-mutually exclusive routes have been proposed. One potential mechanism is dissemination of the virus to ganglia via VZV-infected lymphocytes. Indeed, VZV can infect T cells, and intravenous injection of VZV-infected T cells in a human fetal DRG xenograft SCID-hu mice model resulted in transfer of the virus to the transplanted neurons (9, 10, 29–31). In addition, T cells infected by the closest relative to VZV, simian varicella virus, have been found in the ganglia of primates during primary infection (32). More recently, other immune cells (monocytes, NK cells, NKT cells, and B cells, in addition to both CD4^+^ and CD8^+^ T cells) have been shown to support productive infection of VZV (33, 34) and could potentially migrate and transfer the virus to sensory neurons. Such routes of infection would presumably enable direct access of the virus to the neuronal cell body. Our previous reports using HMN were less supportive for this hematogenous route for VZV latency as HMN are highly susceptible to VZV lytic infection by cell body infection (17) while axonal infection facilitated establishment of latency (16). A non-mutually exclusive proposal by which VZV accesses sensory ganglia is that retrograde axonal transport of the virus occurs from nerve endings innervating the dermis adjacent to cutaneous varicella lesions. This has been supported by the detection of viral antigens in Schwann cells and peripheral nerve axons in patients with varicella, as well as observations that herpes zoster occurs at the site of vaccine inoculation or at sites most affected by primary varicella infection (35, 36). Recent *in vitro* studies have provided direct visual evidence of retrograde axonal transport of VZV (13), and cell-free virus infection by this route also enabled establishment of an *in vitro* latent state using human ESC-derived mixture neurons (15, 16). Intriguingly, we observed that axonal infection of sensory neurons resulted in an *in vitro* latent state more similar to latency in human TG than human ESC-derived mixed neurons, while neuronal cell body infection of HSN was met with marked resistance to lytic infection. Thus, our observations suggest that sensory neuronal latency could occur in a setting in which both hematogenous direct transfer and retrograde axonal transport occur, and further investigation of the interaction between immune cells and sensory neurons is warranted.

The cell bodies of sensory neurons are primarily located in the ganglion. While much has been learned in the past few decades regarding the electrophysiological and molecular characteristics of these sensory neurons, it has only recently been appreciated that other cells and structures within the DRG may contribute substantially to neuronal function (37). Satellite cells, for example, are a specific type of glia that form a close functional relationship with neurons within the DRG and can modify the neuronal microenvironment. Indeed, recent studies have demonstrated a critical role for these cells in the development of pain (38). Other immune cells, including macrophages, T lymphocytes, B lymphocytes, and mast cells are also present within the DRG and may also modulate neuronal function (39, 40). While these other cell types may contribute to VZV neuropathogenesis *in vivo*, the absence of such cells in our *in vitro* model of VZV infection indicates that sensory neurons possess intrinsic, cell-autonomous properties that facilitate viral latency.

We and others have previously reported lytic infection of ESC- and iPSC-derived human sensory neurons, without noting resistance to infection (18, 19). In these reports, co-expression of the nuclear marker Brn3a and cytoplasmic marker peripherin served to mark sensory neurons; however, the maturation state of the cells was unclear as neurons were only differentiated for up to three weeks and additional markers of mature sensory neurons were not assessed. We found that Brn3a and peripherin were expressed early on during differentiation (within two weeks), while additional markers of mature sensory neurons, such Na_v_1.7 and Na_v_1.8, were increasingly expressed following the four week time point of differentiation, consistent with other reports (41). Thus, it is possible that the maturation state of human sensory neurons governs the relative resistance to lytic infection by VZV. Taken together we find that mature human sensory neurons possess intrinsic properties that restrict lytic infection and facilitate the development of VZV latency.

## MATERIALS and METHODS

### Cells

Human iPSC-derived sensory neuron progenitors (HSN; ax0055, Axol Bioscience) were plated on a 24-well plate (1 x 10^5^ cells/well) or a microfluidic platform (7.5 x 10^4^ cells/sector) in Neuronal Plating-XF Medium (Axol Bioscience). Fabrication of a microfluidic platform was previously described (16, 18). Prior to plating the HSN progenitors, a plate or microfluidic platform was coated with poly-L-ornithine (Sigma-Aldrich) (20 µg/mL) or poly-D-lysine (Sigma-Aldrich) (200 µg/mL) in molecular grade water at room temperature overnight, washed with distilled water twice and coated with Matrigel (Corning) (1 µg/mL) in Knockout DMEM/F-12 medium (Thermo Fisher Scientific) for two hours at room temperature following overnight incubation at 37°C in a humified 5% CO_2_ incubator. At one day after plating, the medium was replaced to the complete maintenance medium consisting of Neurobasal Plus Medium, B-27 Plus Supplement (2% [vol/vol]), N2 Supplement (1% [vol/vol]), GlutaMAX-I (2 mM) (Thermo Fisher Scientific), ascorbic acid (200 µM; Sigma-Aldrich), GDNF (25 ng/mL), NGF (25 ng/mL), BDNF (10 ng/mL) and NT-3 (10 ng/mL) (Peprotech) for sensory neuronal maturation. Two days after the plating the HSN progenitors, cells were treated with the complete maintenance medium with mitomycin C (2.5 µg/mL; Nacalai Tesque, Inc) for two hours to eliminate proliferating cells, washed with the complete medium twice and cultured in the complete maintenance medium with replacement of half the volume of culture with fresh media every four days. During maturation in the microfluidic platform, culture medium level in the axonal compartment was kept higher than that in the somal compartment to prevent cell migration to the axonal compartment. H9 human ESC-derived neural stem cells (NSCs) (Passage four to ten) were cultured in proliferating media consisting of Knockout DMEM/F-12 media supplemented with GlutaMAX-I (2 mM), bFGF (20 ng/mL), EGF (20 ng/mL) and StemPro Neural Supplement (2% [vol/vol]) (Thermo Fisher Scientific). NSCs were differentiated into HMNs utilizing a neuronal differentiation medium prepared in Neurobasal Medium with B-27 Serum-Free supplement (2% [vol/vol]) (Thermo Fisher Scientific), and GlutaMAX-I (2 mM). Cells were seeded at a density of 0.5 - 1 × 10^5^ cells/cm^2^, plated in proliferation media for two days and were then differentiated for four to six weeks before experiments were performed. Human retinal pigmented epithelium ARPE-19 cells (American Type Culture Collection [ATCC] CRL-2302) were maintained in DMEM/F-12+GlutaMAX-I (Thermo Fisher Scientific) supplemented with heat-inactivated 8% FBS (fetal bovine serum; Sigma-Aldrich).

### Immunostaining and imaging

HSNs and HMNs were washed once with phosphate-buffered saline (PBS) and fixed with 4% paraformaldehyde for 20 minutes (min) at room temperature. Cells were washed with PBS before treatment with 0.25% Triton X-100 and 5% normal donkey serum for one hour. Primary antibodies, mouse anti-Na_v_1.7 monoclonal antibody (clone N68/6, Abcam) (1:100), mouse anti-Na_v_1.8 monoclonal antibody (clone N134/12, Abcam) (1:100), goat anti-Peripherin polyclonal antibody (C-19, Santa Cruz) (1:200), mouse anti-Brn3a monoclonal antibody (clone 5A3.2, Chemicon) (1:100), chicken anti-Neurofilament polyclonal antibody (NFM, Aves Lab) (1:200), mouse anti-Islet 1 monoclonal antibody (clone 1B1, Abcam) (1:200), rabbit anti-Tuj1 polyclonal antibody (Poly18020, Covance) (1:200),and mouse anti-Tuj1 monoclonal antibody (clone TuJ-1, R&D systems) (1:500) were used to stain overnight at 4°C. After three washes with 1X PBS, appropriate Alexa Fluor 488/555/647-conjugated anti-rat/rabbit/mouse/goat/chicken secondary (1:250, Thermo Fisher Scientific) in donkey serum were incubated for 1.5 hours (hrs) at room temperature. Finally samples were counter stained for nucleus with 1 μM DAPI (4′, 6-diamidino-2′-phenylindoldihydrochloride; Thermo Fisher Scientific). Zeiss inverted Axio Observer fluorescent microscope (Zeiss, Germany) was used to image the cells.

### Viral infections

Cell-free virus of VZV strain pOka (parental Oka) or rVZV_LUC_BAC (derived from pOka) reconstituted in MRC-5 cells by transfection of VZV_LUC_BAC DNA (from Hua Zhu, New Jersey Medical School, Rutgers University, Newark, NJ) were prepared and titrated as described previously (18, 42).

For lytic infection, neurons were infected with cell-free virus for two hours in 300 µL medium, washed with the medium twice, treated with low pH buffer (40 mM sodium citrate, 10 mM potassium chloride, 135 mM sodium chloride [pH 3.2]) for 30 seconds (sec), washed with the media once and cultured for indicated durations. For VZV *in vitro* latency, we applied slight modifications to our previous methodology (16, 18). Briefly, neurons were differentiated in a microfluidic platform for 54 days and infected from axonal compartment with 10 µL of the cell-free virus (400 pfu titrated on ARPE-19 cells) with 10 µL media. After two hours infection, inoculum was removed, and axonal compartments were treated with the low pH buffer for 30 sec, washed with the media and cultured for two weeks.

To visualize infectious foci on ARPE-19 cells, cells were fixed with 4% paraformaldehyde/PBS (Nacalai Tesque, Inc.), stained with mouse anti-gE monoclonal Ab (clone 9) (1:10 dilution in PBS) (43), followed by anti-mouse IgG horseradish peroxidase (HRP)-linked whole Ab sheep (1:5,000 dilution in PBS) (GE Healthcare Bio-Sciences), and reacted with 3, 3’, 5, 5’-tetramethylbenzidine-H peroxidase substrate (Moss, Inc.).

### Western Blot

Proteins were harvested four to five days post virus infection in RIPA Lysis and Extraction Buffer (Boston Bio) with 1X Protease Inhibitor and 1X Phosphatase Inhibitor (Thermo Fisher Scientific). Protein Concentration was determined using BCA Assay Kit (Thermo Fisher Scientific) as per manufacturer’s protocol. 20 µg of protein were separated by loading in 4-15% MINI-PROTEAN TGX gel (Bio-Rad) followed by transfer onto 0.2 µm nitrocellulose membrane, Trans-Blot Turbo pack (Bio-Rad). Membrane was further blocked with 5% milk in 1X PBS and Tween 20 for 30 min followed by incubation with mouse anti-gE monoclonal antibody (clone 8612, EMD Millipore) (1:3,000) and appropriate control rabbit anti-GAPDH monoclonal antibody (clone D16H11, Cell Signaling Technology) (1:5,000) overnight at 4°C. The following day, the membrane was probed with HRP-conjugated anti-mouse or anti-rabbit (1:5,000, GE Healthcare Bio-Sciences) secondary antibody after washing several times with 1X PBS for 45 min at room temperature. The antibody binding was detected using SuperSignal West Femto (Thermo Fisher Scientific) incubated for five minutes in dark and visualized using Universal Hood II Gel Doc System (Bio-Rad).

### Flow cytometry

Cells were washed once after removing culture media with 1X PBS and then incubated in Accutase (Sigma-Aldrich) for 10 min to harvest the cells. The cells were then neutralized and centrifuged at 200 x g for four minutes to collect pellets. The samples were then re-suspended in 500 μL of cold 1X PBS and transferred into 5 mL polystyrene round bottom flow tube with cell strainer cap (Corning). Samples were then analyzed for GFP positive cells for each time point and condition until the cell count events reached at least 10,000.

### Nucleotide extraction and Quantitative PCR

DNA and RNA from VZV-infected human sensory neurons were isolated using the FavorPrep Blood/Cultured Cell Total RNA Mini Kit (Favorgen Biotech) in combination with the NucleoSpin RNA/DNA buffer set (Macherey-Nagel). DNA was first eluted from the column in 100 µL DNA elution buffer, the column was treated with recombinant DNase I (20 units/100 µL; Roche Diagnostics) for 30 min at 37°C and RNA was eluted in 50 µL nuclease free water. RNA was directly treated with Baseline-ZERO DNase (2.5 units/50 µL; Epicentre) for 30 min at 37°C. cDNA was synthesized with 12 µL of RNA and anchored oligo(dT)_18_ primer in a 20 µL reaction using the Transcriptor First Strand cDNA synthesis kit at 55°C for 30 min for reverse transcriptase reaction (Roche Diagnostics).

DNA or cDNAs were subjected to quantitative PCR (qPCR) using KOD SYBR qPCR Mix (TOYOBO) in the StepOnePlus Real-time PCR system (Thermo Fisher Scientific) (1 µL of DNA or cDNA per 10 µL reaction in duplicate). All primer sets used for qPCR (**Table 1**) were first confirmed for the amplification rate (98-100%) using 10-10^6^ copies (10-fold dilution) of pOka-BAC genome or VLT plasmid (21) and the lack of non-specific amplification using water. The qPCR program is as follows; 95°C for 2 min (1 cycle), 95°C for 10 sec and 60°C 15 sec (40 cycles), and 60 to 95°C for a dissociation curve analysis. Data is presented as relative VZV level to cellular beta-actin (cDNA) or CD24 (DNA) defined as 2^-(Ct-value VZV gene - Ct-value beta-actin or CD24)^.

**Table 1.**
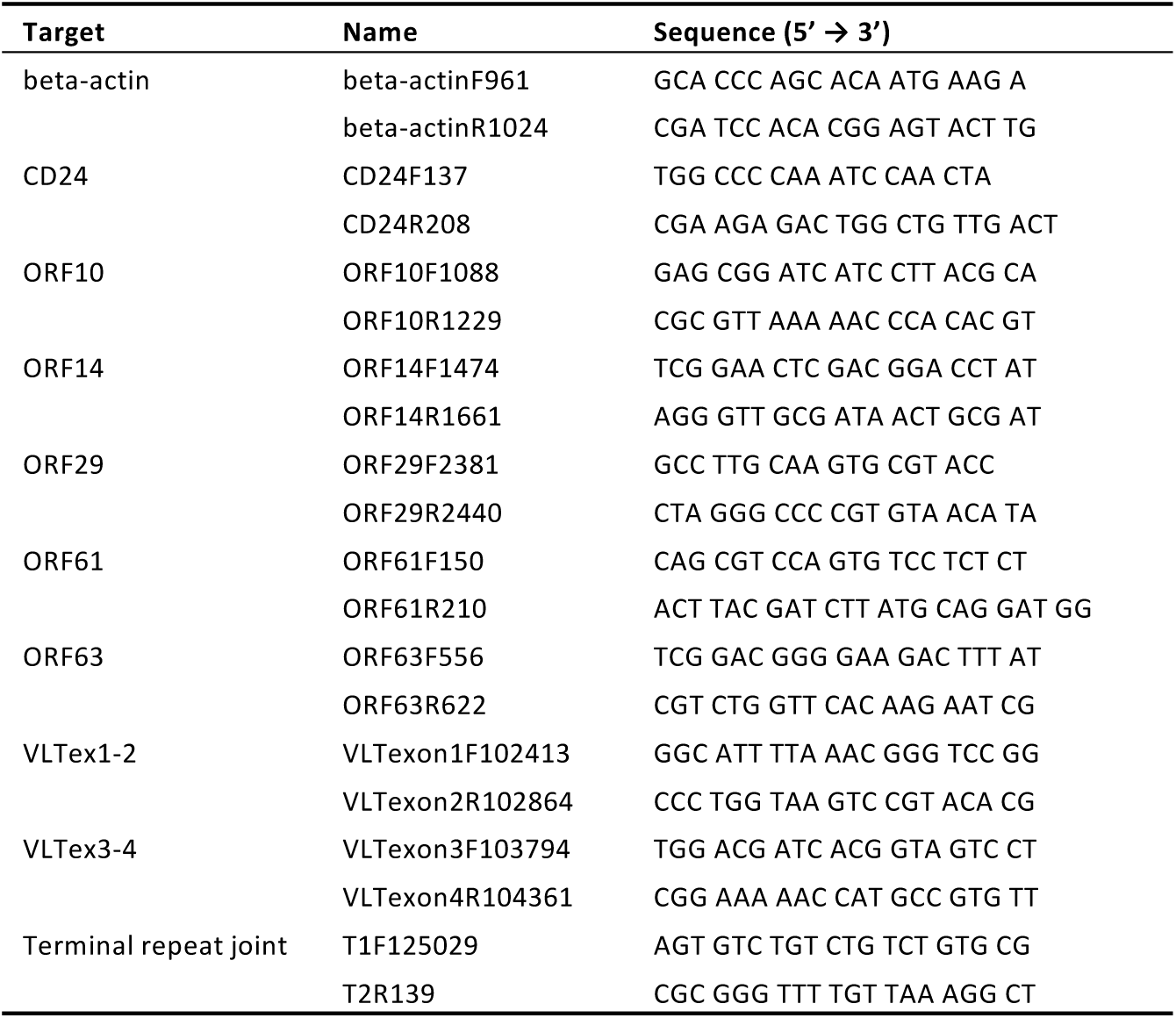
Primers for qPCR assay.

## ACKNOWLEDGEMENTS

This work was supported by the National Institutes of Health (R21 NS107991 to A.V.) and the Takeda Science Foundation, Daiichi Sankyo Foundation of Life Science, Japan Society for the Promotion of Science (JSPS KAKENHI JP17K008858, JP16H06429 and JP16K21723) and the Ministry of Education, Culture, Sports, Science and Technology (MEXT KAKENHI JP17H05816) (T. S.).

